# Tracing co-regulatory network dynamics in noisy, single-cell transcriptome trajectories

**DOI:** 10.1101/070151

**Authors:** Pablo Cordero, Joshua M. Stuart

**Affiliations:** UC Santa Cruz Genomics Institute, University of California, Santa Cruz, California, USA

**Keywords:** single-cell measurements, Gaussian mixtures, transcriptomics, single-cell trajectory reconstruction

## Abstract

The availability of gene expression data at the single cell level makes it possible to probe the molecular underpinnings of complex biological processes such as differentiation and oncogenesis. Promising new methods have emerged for reconstructing a progression ‘trajectory’ from static single-cell transcriptome measurements. However, it remains unclear how to adequately model the appreciable level of noise in these data to elucidate gene regulatory network rewiring. Here, we present a framework called Single Cell Inference of MorphIng Trajectories and their Associated Regulation (SCIMITAR) that infers progressions from static single-cell transcriptomes by employing a continuous parametrization of Gaussian mixtures in high-dimensional curves. SCIMITAR yields rich models from the data that highlight genes with expression and co-expression patterns that are associated with the inferred progression. Further, SCIMITAR extracts regulatory states from the implicated trajectory-evolving co-expression networks. We benchmark the method on simulated data to show that it yields accurate cell ordering and gene network inferences. Applied to the interpretation of a single-cell human fetal neuron dataset, SCIMITAR finds progression-associated genes in cornerstone neural differentiation pathways missed by standard differential expression tests. Finally, by leveraging the rewiring of gene-gene co-expression relations across the progression, the method reveals the rise and fall of co-regulatory states and trajectory-dependent gene modules. These analyses implicate new transcription factors in neural differentiation including putative co-factors for the multi-functional NFAT pathway.

## Introduction

Understanding the dynamics of gene expression progression in a cell population as it traverses a biological process such as differentiation has been an outstanding problem in modern cell biology. These dynamics are characterized not only by the changes in cell-to-cell gene expression levels, but by the rewiring of gene regulatory networks as the cells transform from one transcriptional state to another. Tracking these gene regulatory changes would pinpoint coordination of biological function as gene modules are turned on or off throughout the progression.

Single-cell transcriptomics has given important insights into gene expression dynamics, revealing the stochastic nature of gene expression and characterizing in detail the behavior of small genetic networks.^1–4^ In their initial incarnation, these measurements were confined to demanding microscopy protocols that assayed gene expression levels through time of only a handful of genes. In recent years, advances in flow cytometry, microfluidics, and sequencing technologies have enabled the interrogation of up to the whole transcriptome in hundreds to thousands of cells.^5–7^ Application of these techniques to biological processes such as development provide snapshots of cell states through time and space.

Many computational methods have emerged to infer trajectories of connected state transitions from the static samplings of single-cell transcriptomes. The goal of these methods is to provide a pseudotemporal ordering of cells in which neighboring cells are similar to each other, capturing an overall biological progression. These approaches have been successfully applied to elucidate complex transcriptional patterns and regulators in myoblast differentiation,^8^ B cell development,^9^ and haematopoiesis.^10^ Nevertheless, cell orderings alone give little insight into the state of gene regulatory networks across time. In addition, while most methods use strategies to tackle biological and technical noise, none account for the dynamic, heteroscedastic nature of the data. Further, only a few take into consideration uncertainties in pseudotime assignments,^11^ making error estimates difficult to evaluate.

To address these challenges we propose a strategy, Single Cell Inference of MorphIng Trajectories and their Associated Regulation (SCIMITAR), for inferring gene expression network dynamics throughout biological progression from static, single-cell transcriptomes. SCIMITAR gives a detailed, fully probabilistic description of the expression trajectory that, in contrast with previous methods, explicitly accounts for heteroscedastic noise in the data. In addition, it tracks the changes of gene-gene expression correlations at each point in the progression. The probabilistic nature of SCIMITAR transition models allows for evaluating the shape of the multivariate gene expression distribution as a function of biological progression, which we show can be used to pinpoint co-regulatory cell states.

We benchmarked SCIMITAR’s inference capabilities in two scenarios. First, we tested its ability to infer cell ordering and network rewiring from simulated transcriptomic measurements where the underlying cell behavior was known. Second, we asked whether SCIMITAR could yield insights in the developmental trajectory of human fetal neurons by analyzing recently published fetal brain single-cell measurements. A likelihood ratio test designed for SCIMITAR revealed 36 genes that significantly varied throughout the progression but that were missed by standard differential expression between cell groups including genes in cornerstone developmental pathways such as the hypoxia inducible factor 1 *α* (HIF1*α*), nuclear factor of activated T cells (NFAT), and androgen receptor (AR) pathways. Further, by tracking SCIMITAR co-expression matrices across pseudotime we were able to detect the evolution of co-regulatory states, gene modules, and genes that gained and lost connectivity throughout the trajectory.

## Results

### Uncovering the full probability distribution progression underlying static single-cell measurements with SCIMITAR

Recently, there has been an explosion of single-cell transcriptomic data in various biomedical contexts and systems. A projection of the data from three such studies (refs^8,10,12^) in Fig 1A using a locally linear embedding reveals that these datasets are characterized by distinct groups of many cells interspersed with cells that fall along what appear to be isolines between groups. This structure suggests a model that combines distributions for cell population density and evolving cell states with heteroscedastic noise. One such model that could describe these data is a continuous mixture of Gaussian distributions with constraints that allow only for smooth, continuous changes in parameters over the course of the progression. We call such a model a Morphing Gaussian Mixture (MGM, see Methods and Fig 1B). The MGM has a mean function, *µ*: [0, 1] → ℝ^*n*^ that threads through the data and is equipped with a covariance matrix function Σ: [0, 1] → ℝ^*n*×*n*^ that defines a Gaussian distribution at each point in the progression, with *n* being the number of genes. The mean and covariance matrix functions vary continuously throughout the [0, 1] interval, defining a probability *P* (*x*|*µ*, Σ, *t*) for each cell gene expression vector *x* and pseudo time-point *t* ∈ [0, 1]. To ease inference, these mean and covariance functions can be parametrized with different functional classes, such as polynomials, splines, or Gaussian processes (see Methods). This probabilistic structure maps samples to a smooth curve and allows points to veer away stochastically by modeling the structure of the changing biological and technical noise. *P* (*x*|*µ*, Σ, *t*) captures the uncertainty of a cell mapping to a particular pseudotime due to the changing covariance nature of the MGM. A key advantage of this approach is that it replaces standard, grouped differential gene expression analysis or differential co-expression analysis with a more sensitive test for potential gene-gene regulatory relationships that change throughout the progression. Details of the MGM model as well as inference of its parameters from data are given in the Methods section.

**Fig. 1.**
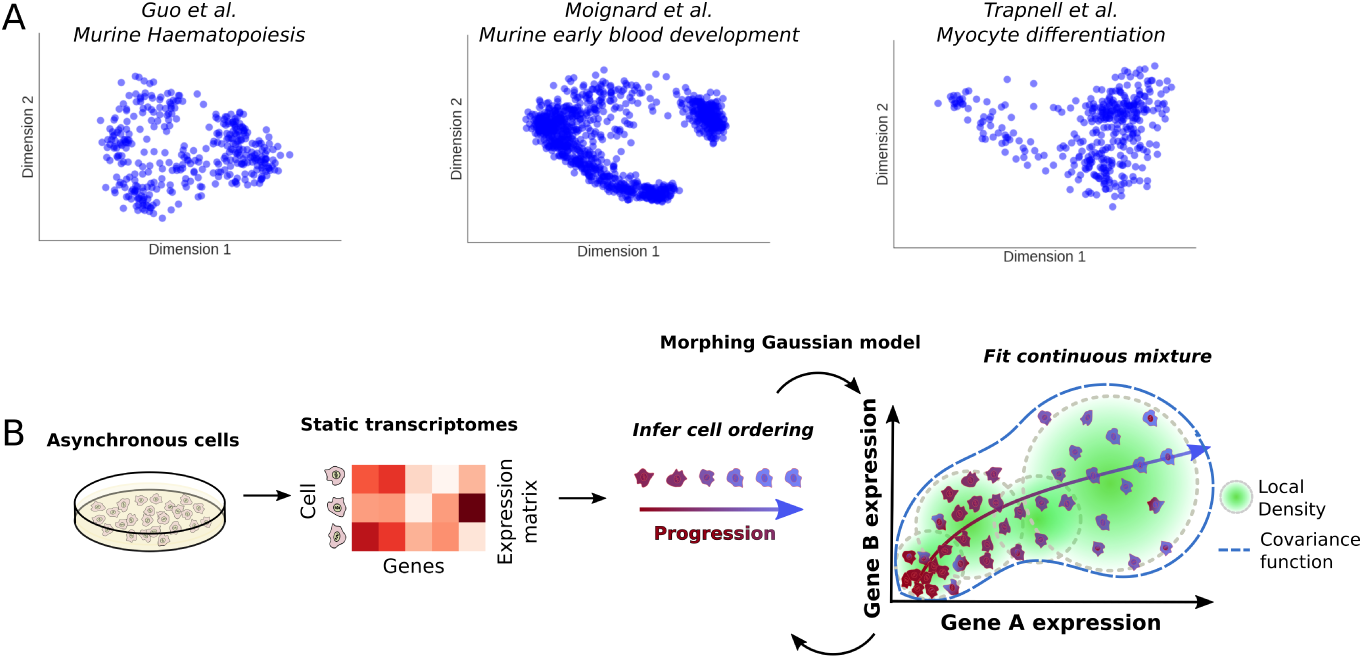
A. Survey of three different single-cell transcriptomic studies. From left to right: murine haematopoiesis by Guo et al., early blood development by Moignard et al., and myocyte differentiation by Trapnell et al. B. Overview of the SCIMITAR method. Trajectory modeling with dynamic and correlated noise of static transcriptomes of asynchronous cells is achieved by iterating through optimal cell ordering and inference of a continuous set of Gaussian distributions in a morphing mixture of Gaussian models (see Methods in text).

### Benchmarking SCIMITAR in simulated data

To test our strategy, we asked whether SCIMITAR could infer the underlying cell ordering and co-expression networks of simulated data where the ground truth was available. We tested SCIMITAR’s cell order inference capabilities in two settings in which noise was added to the system: 1) the noise is *uncorrelated* to the underlying trajectory and 2) the noise is *correlated* with the trajectory. The first setting, adding noise uncorrelated with the trajectory, tests robustness of the method in the presence of genes that are unrelated to the biological progression and that confound ordering inference. The second setting tests how biological and technical noise intrinsic to the system, including gene-gene correlated noise that change over time, affect cell ordering inference.

For the first setting, we simulated data closely following the simulation procedure described in ref.^9^ We simulated data in which 3 genes defined the true cell state and 7 genes represented unrelated (uncorrelated) expression programs to the simulated progression. Simulations in this scenario then, 3 dimensions of the data were “signal” while 7 were “noise”. To obtain the three-dimensional trajectory, we performed a random walk for 600 steps and sampled a ‘cell’ from a standardized normal distribution centered at the current point in the walk. We then added seven dimensions of Gaussian noise. We generated several datasets with an increasing noise magnitude (quantified as the standard deviation times the range of the trajectory). We then used SCIMITAR to model these data and obtain the model’s optimal cell ordering. We used SCIMITAR with three different functional classes (see Methods): third degree polynomials, cubic splines, and Gaussian Processes with a squared exponential correlation function (GP). We compared SCIMITAR’s performance with the cell orderings inferred by two popular methods, Monocle^8^ and Wanderlust,^9^ and used the Pearson correlation coefficient to compare the approaches (see Fig 2A). The best overall performers were all SCIMITAR models, with Wanderlust coming in close second and Monocle performing slightly worse possibly due to its assumption of linearity in its dimensionality reduction step in agreement with previous studies.^13^ All methods were susceptible to the noisy dimensions uncorrelated with the trajectory.

**Fig. 2.**
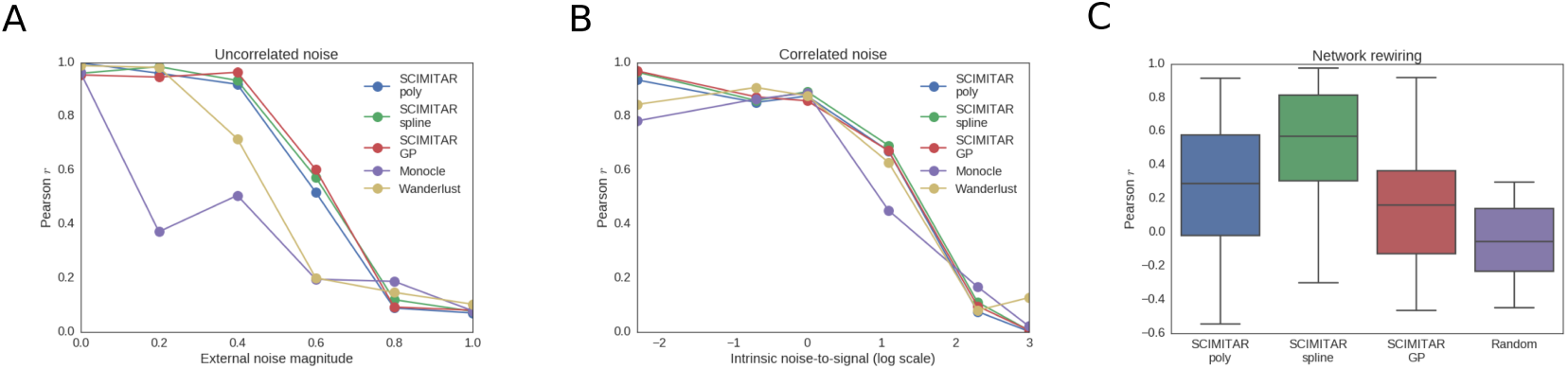
SCIMITAR *in silico* benchmark. A. Cell ordering results for three functional classes of SCIMITAR (a third degree polynomial, a cubic spline, and Gaussian processes with squared exponential correlation model) and two state-of-the-art methods Monocle and Wanderlust in a setting with noise uncorrelated to the trajectory. B. Cell ordering results for noise correlated with the trajectory. C. Evaluation results of network rewiring across biological progression for SCIMITAR’s three functional classes and random covariance functions.

For the second test that adds noise correlated with the trajectory, we simulated a curve, *µ*_*sim*_ traversing a 10-dimensional space using 10 randomly-generated quadratic polynomials. The correlated noise was simulated from the evolution of randomly generated Watts-Strogatz networks and an additional set of quadratic polynomials with 6 different settings of signal-to-noise ratios (see Supplemental Methods for a detailed description of this benchmark). We found all methods performed similarly (Fig 2B), suggesting that noise intrinsic to the system, including gene-gene statistical dependencies, equally confounds any cell ordering inference method.

In addition to solving the cell ordering problem, SCIMITAR models track evolving gene-gene correlations. We used the correlated noise simulations to test the accuracy of SCIMITAR’s gene network rewiring inference. To this end, we compared the covariance functions inferred by the polynomial, spline, and GP SCIMITAR versions. We measured the concordance of trends between each entry of the predicted matrix functions 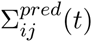 and the corresponding entry of the simulated values 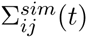 using the Pearson correlation coefficient (see Fig 2C). The spline version of SCIMITAR produced the highest correlation coefficients while all versions were substantially better than randomly-generated covariance matrix functions. Closer examination of the three functional classes revealed that the GP version tended to overfit the data locally, closely following local covariance structure even in regions where a few samples were present while the polynomial version lacked the flexibility to model some complex twists and turns in evolving true covariance structures. The spline version struck a balance between smoothing inferences in intervals of the trajectory with few samples and maintaining flexibility to capture non-linear trends. We therefore chose to use the spline functional class for SCIMITAR models in the remainder of this study.

### A differentiation model for human fetal neurons

In a previous study, Darmanis et al. obtained a transcriptomic map of the adult and fetal brain using single-cell RNA-seq measurements.^14^ One of the findings of the study was a continuous transition the between fetal replicating and quiescent neurons. We applied SCIMITAR to infer cell ordering and network rewiring of these data to elucidate key regulatory changes across the differentiation process. We downloaded these data from the gene expression omnibus (series identifier GSE67835) and obtained the subset corresponding to all fetal neurons. We focused on all transcription factors that were expressed in at least 10% of the cells, log-transformed the data and controlled for cell-cycle effects using scLVM.^15^ We then fit SCIMITAR to the data and visualized the results in a two-dimensional locally linear embedding (see Fig 3A). The visualization suggested a single linear trajectory that traversed the fetal replicating and quiescent neurons which was captured by the SCIMITAR model. To obtain progression associated genes, we used a likelihood ratio test tailored for SCIMITAR models with dynamic noise (see Methods). The test revealed 92 genes with expression that was significantly psuedotemporal-dependent (see Fig 3B). To obtain global insights from these genes, we used hierarchical clustering with the Pearson correlation similarity metric to group them into 5 groups and performed Gene Ontology and KEGG pathway enrichment tests on each group (see color groups in Fig 3B). Early-expressed genes (red and green clusters) were associated with glucocorticoid receptors, heat shock factors, and signal transduction; genes expressed in the middle of the progression (yellow and pink clusters) were enriched with Maf-like proteins and cytokines; and the late-expressed genes (cyan cluster) had apoptosis, neurogenesis, and alternative splicing enrichment. These enrichments correspond to multiple observations in the literature. For example, heat shock factor proteins are well known to be involved in early neurodifferentiation^16^ while glucocorticoid receptors and Maf-like proteins are found to be expressed at different stages in hippocampal and developmental neurogenesis, respectively.^17,18^ Further, neurodifferentiation has been found to be particularly enriched for alternative splicing events.^19^

**Fig. 3.**
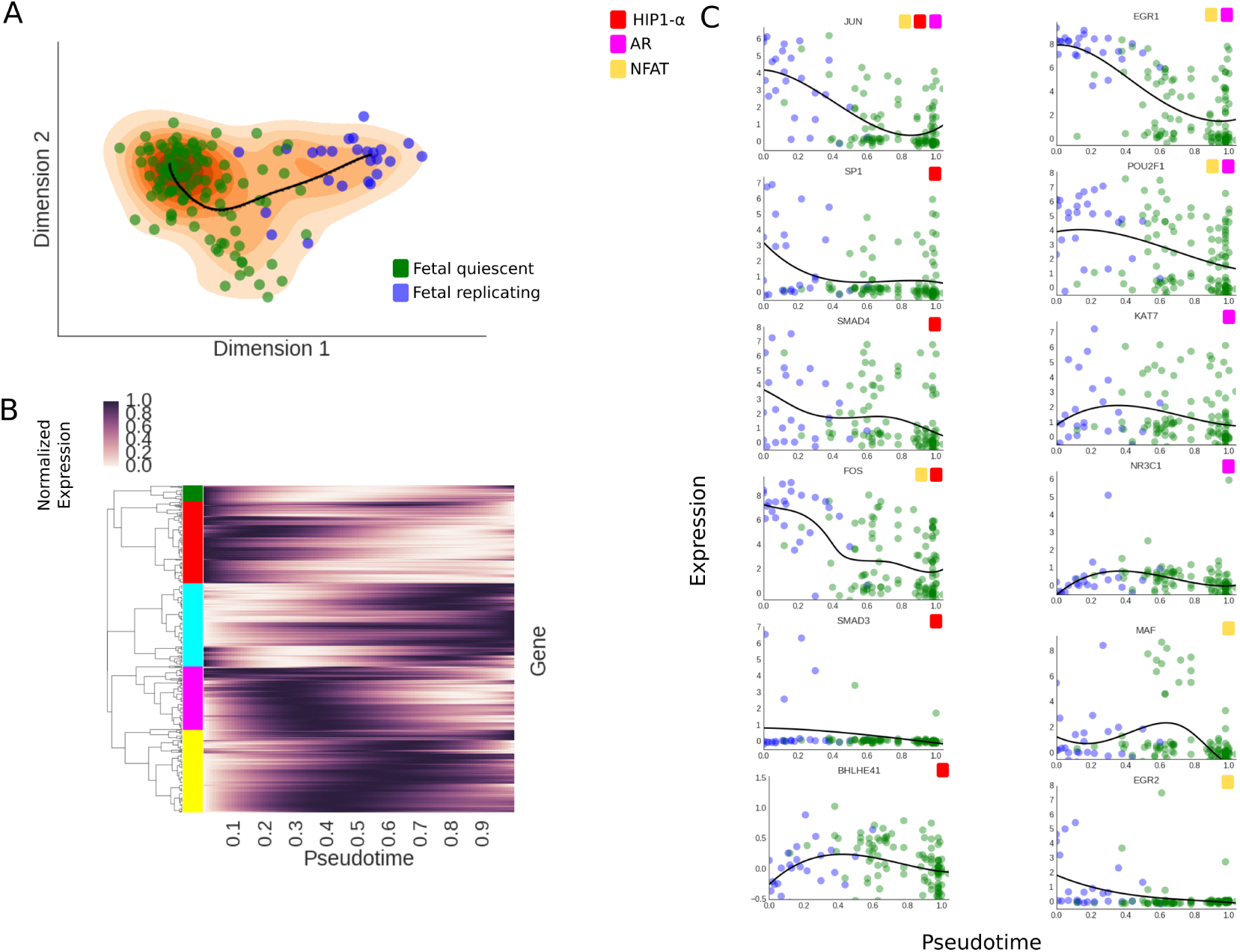
A. SCIMITAR model for fetal neuron differentiation, projected to a 2-dimensional locally linear embedding. The data is plotted as circles in blue (fetal replicating neurons) and green (fetal quiescent nuerons) while the SCIMITAR model’s mean is plotted in black and its projected PDF is plotted in orange. B. Normalized SCIMITAR model means for genes that were deemed progression associated across the progression, clustered into five different clusters using expression correlation throughout psuedotime. C. Expression levels of several genes from three central neurodifferentiation pathways: the HIF1*α*, NFAT, and Androgen Receptor (AR) pathways that were pinpointed by SCIMITAR associated progression tests.

We then compared SCIMITAR’s progression associated genes to those obtained using an ANOVA differential expression test between cells grouped according to their fetal replicating or quiescent annotations. SCIMITAR uncovered 36 genes missed by ANOVA, most of which were highly expressed in the middle of the progression, a detail that is lost when grouping cells into two groups. These missed genes implicate different pathways whose genes were engaged in progression dynamics. For example, five genes, BHLHE40, SMAD3, SP1, and SMAD4, of the hypoxia inducible factor 1 *α* (HIF1*α*) pathway, involved in neural development,^20^ were revealed to follow an ordered progression by the SCIMITAR model but missed using grouped ANOVA differential expression (see Fig 3C). SCIMITAR revealed that the progression associated genes of this pathway were mostly active in early stages of differentiation. SCIMITAR also illuminated two other pathways: the Nuclear factor of activated T-cells (NFAT) and the Androgen receptor pathway which is critical for neural stem cell fate commitment^21,22^ (see Fig 3C).

We note that SCIMITAR’s progression associated genes did not include 7 genes from the ANOVA list, false positives for which the variance was too large or where the statistic was skewed by outliers in an otherwise lowly expressed gene. Nevertheless, three genes that seem to be be differentially expressed by manual inspection (BCL11B, AFF1, and REST) were found by ANOVA but missed by SCIMITAR, presumably due to a small subset of cells driving the change between groups.

### Evolving co-expression networks reveal defined co-regulatory states

We then used SCIMITAR’s inferred covariance functions to track changes in gene-gene connectivity across the progression. We sampled 100 correlation matrices at regular intervals from the covariance function, restricting the matrices to genes deemed progression associated. We calculated a global distance matrix between networks using Frobenius distance to assess their similarities and plotted the similarity values across pseudotime (see Fig 4A). As expected, the strongest similarities were between networks that were neighbors in pseudotime. However, three network clusters could be appreciated in the matrix, suggesting three different co-regulatory states. We obtained the consensus network of each state by averaging the network members of the cluster. Then, we ranked each gene by comparing their co-expression degree in each state to their co-expression degrees in the other two states using z-scores. The top 20 genes that gained the most connectivity in each state are listed in Fig 4B. All of the gain-of-connectivity genes include genes that have been established as key players in neurodifferentiation, such as PAX6, DLX1, and NEUROD6 and were enriched with neurodevelopmental and neurogenesis GO terms.

**Fig. 4.**
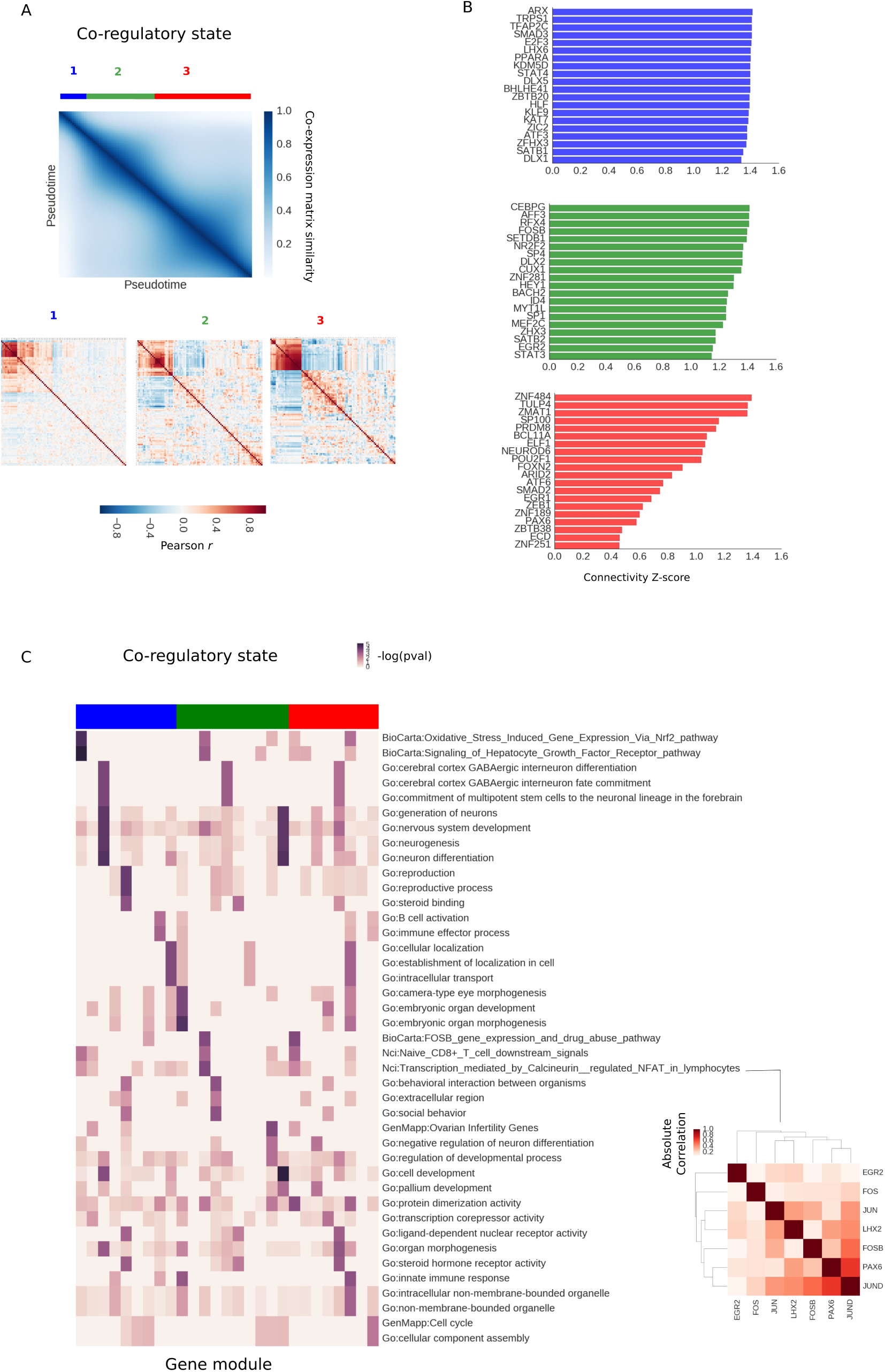
A. Similarity matrix between co-expression matrices fitted in the SCIMITAR fetal neuron differentiation model across pseudotime. Three different co-regulatory states can be appreciated in the matrix, marked in blue, green, and red. B. Top 20 genes with the most gain-of-connectivity in each co-regulatory state alongside their log co-expression degree. C. Evolution of annotated modules. Each column is a module and each row is a gene annotation — enrichments are shown as *−log*(*p − value*) in the heatmap. Column colors denote co-regulatory state. An NFAT-associated module of state 2 is highlighted in the red matrix

To track highly connected gene modules of each state that significantly changed their connectivity, we obtained gene modules for each co-regulatory state using affinity propagation (with a dampening parameter of 0.5), finding 27 gene modules in total. We annotated these modules by gene set enrichment and ordered them across pseudotime (see Fig 4C). This analysis revealed a coordinated functional response across the trajectory: modules in state 1 were annotated with neural stem cell commitment, immune response, and protein trafficking, while state 2 was enriched with embryonic development, neuron regulation, and pallium development. State 3 had more diverse enrichments, from morphogenesis to membrane organelles, suggesting a stage when cells start taking on mature neuron roles depleted of differentiation potential. Importantly, this analysis pinpointed an NFAT-associated module to be most active in co-regulatory state 2 (see Fig 4D). Most NFAT co-factors involved in neural development are still unknown.^23^ The uncovered NFAT-associated module provides putative candidates for this function. The full list of modules and their gene networks can be found in the Supplemental Results (see below).

## Discussion

An outstanding goal of systems biology is to understand the principles under which the gene regulatory circuitry of a cell changes during a biological process. Single-cell transcriptomes offer a fast way to obtain transcriptome-wide snapshots of these processes. When properly analyzed, these data can be used to recover the principal trends of the biological progression, but current methods do not model the dynamic gene-to-gene correlations in expression that are the hallmarks of the underlying regulatory circuitry. Here, we presented SCIMITAR, a strategy that leverages morphing Gaussian mixtures to track biological progression and model the rewiring of these gene networks from static transcriptomes. SCIMITAR models account for heteroscedastic noise and increase the statistical power to detect progression-associated genes when compared to traditional differential expression tests. Further, the models allow for detecting modes in co-expression structure in the trajectory: defined co-regulatory states that represent potential metastable and transitionary cell states. We note that Gaussian mixtures with non-diagonal covariance matrices suffer from the curse of dimensionality, which we have tried to control for by using shrinkage estimators. Exploring the robustness of other types of regularized estimators such as the graphical LASSO would be a logical next step to improve confidence in the inferred morphing mixture models.

SCIMITAR is part of a recent wave of probabilistic methods for cellular trajectory reconstruction from single-cell measurements.^11,24^ These types of models present several advantages, such as assigning uncertainty estimates of cell orderings and providing a natural way for mapping new samples to a trained model — a necessary task for building queryable trajectory maps with multiple progressions. Although SCIMITAR as presented cannot model branched cellular trajectories such as those corresponding to multiple cell fate decisions, the framework can be readily extended by replacing the single-curve parametrization of the mixtures with a branching structure, which deserves further investigation.

## Methods

### Morphing Gaussian Mixtures: correlated gene progression modeling with no dimensionality reduction

Single-cell transcriptomic measurements are high-dimensional, with the number of variables measured typically ranging from a few markers (generally no less than 48) to the full transcriptome that can be upwards around 30000 transcripts. However, not every gene or transcript is relevant to the biological system of interest and most are not expressed at all. Further, due to the underlying gene regulatory networks, the expression patterns of many genes are correlated and the strength of this correlation changes throughout the progression as the regulatory system changes from one cell state to the next. These biological constraints put the data in some low-dimensional manifold, a property that is used in various ways by cell ordering algorithms to justify reducing the dimensionality of the dataset to a manageable number of dimensions. Monocle, for example, reduces the data’s dimensionality to 2 dimensions using independent component analysis and performs its calculations on a lower dimensional manifold. While the procedure captures general aspects of the trajectory, 2 dimensions is generally not enough to capture all of the relevant variability of the data and the reduction leads to loss of information that can impact trajectory reconstruction (see e.g. our benchmarks in the Results sections and other benchmarks in^13,24^). Other methods, such as Wanderlust, reduce the dimensionality in a more principled way through nearest-neighbor calculations but forego capturing the changes in gene-gene expression correlations over time. To address both of these shortcomings, we introduce a model that retains the dimensionality of the dataset and tracks gene-gene correlations throughout the trajectory. To this end, we extended Gaussian graphical models to accommodate time-dependent changes in the mean and covariances of the model with time being a latent variable.

Gaussian graphical models are one of the dominant frameworks for analyzing gene expression data, where the data is assumed to follow a multivariate Gaussian distribution defined by a mean vector and a covariance matrix. Modeling the data becomes more challenging in the presence of population structure where several different populations, each with its own distribution, are intermixed. Gaussian mixture models, which posit that the data comes from a finite combination of multivariate Gaussians, have been used successfully in this scenario.^25^ In static single-cell expression from a group of cells continuously undergoing a biological process, such as differentiation, the boundaries between populations are blurred and the data is best described as a continuous transformation between the first and last states. We model this transformation by assuming that the data comes from a *continuous* Gaussian mixture, parametrized by timepoints within the progression (the so-called pseudotime), which are unknown. Let *X* be the data, *p* the number of genes, *µ*: [0, 1] → ℝ^*p*^, Σ: [0, 1] → ℝ^*p*×*p*^ the mean and covariance functions of the evolving populations that are time dependent, and *γ* a probability distribution on the [0, 1] interval representing cell population density at each pseudo time-point. Then the probability of the data given the model *M* = {*µ*, Σ, *γ*} can be written as:

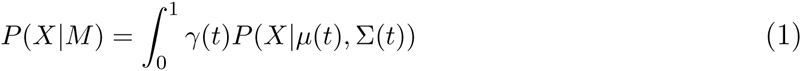

Here, *t* stands for the pseudotime in the progression. This model, which we name the morphing Gaussian mixture model (MGM), differs from other mixture models in that we require the mean and covariance structures to be described through continuous functions and generalize other related models such as principal curves by inferring local covariance structure in addition to the mean curve. The changing covariance structure allows the model to both keep the dimensionality of the dataset and track co-expression changes throughout the progression.

To fit the model to the data, we use a maximum likelihood approach. As previously defined, the parameters in the MGM model are difficult to infer, since optimization of the likelihood function requires searching the space of all continuous functions. Additionally, the positive-definite requirement on Σ(*t*) makes fitting the matrix function difficult. Therefore, we recast the problem of fitting Σ(*t*) into fitting its pseudotime-dependant Cholesky decompositions: Σ(*t*) = *C*(*t*)^*T*^ *C*(*t*), ∀*t* and impose a functional form to the *µ*(*t*) and *C*(*t*) functions. We consider three different functional classes: polynomials, Gaussian processes with squared exponential correlation models, and cubic, De Boor smoothing splines, a special case of Gaussian processes.

To fit the parameters of the model, we employ coordinate ascent. In the first step, we are given a fixed set values for *M* and we calculate, for each sample *x*, the optimal pseudotime *t*_*opt*_ in the [0, 1] interval for which *P*(*x*|*µ*(*t*_*opt*_), Σ(*t*_*opt*_)) is maximized. In the second step, given optimal pseudotime values, we calculate the cell density *γ* by fitting kernel density estimator to the assigned pseudo time-points. Finally, in the third step, given density weights *γ* and pseudotime assignments, we find the *µ* and Σ functions that best fit the data. To achieve this, we approximate *µ*(*t*) and *C*(*t*) locally by obtaining optimal values at the pseudo time-points 0, 0.1, 0.2, …, 1.0, inferring the local mean and covariance using each data point weighted by their probabilities as given by *γ*, and leveraging these values to fit functions from the desired functional class (e.g. a polynomial, spline, or Gaussian process). Because we may have considerably less samples than genes, we use the Ledoit-Wolf-type estimator in the R corpcor package to fit the covariance at each pseudo time-point. We repeat this procedure until convergence, as evaluated by the Pearson correlation coefficient of current and past pseudotimes, with stopping criterion *r* > 0.9. As initial values for pseudotime assignments to our optimization routine, we use a de-noised one-dimensional locally linear embedding.^26^

### Visualization of the data and SCIMITAR models

To visualize the data and models, we use 2-dimensional locally-linear embeddings, with number of neighbors set to 80% of the number of samples. We plot SCIMITAR means by sampling 100 equidistant points across the mean function and projecting to the embedding. To obtain a projection of the SCIMITAR model’s probability density function, we obtain 1000 samples from the model, evenly spaced across pseudotimes in the [0, 1] interval, project to the embedding, and plot a 2-dimensional kernel density estimator of the 1000 points.

### A progression association statistical test

To obtain genes whose expression is progression-dependent, we use a likelihood ratio test to compare the SCIMITAR model of each gene’s progression and the null hypothesis where the expression of the gene is ‘flat-lined’, i.e. does not track with the model’s path. Specifically, we calculate the statistic:

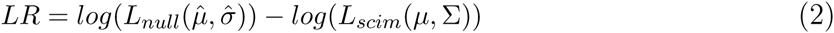

Where *L*_*scim*_, *L*_*null*_ are the likelihood functions of the SCIMITAR and null models, respectively, with the null distribution defined as a normal distribution centered at the empirical mean 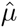 and standard deviation 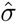 of all the data representing the case where the data is independent of the progression. To assess whether the null hypothesis should be rejected, we obtain the distribution of *LR* under the null hypothesis using parametric bootstrapping with 1000 samples and compare the resulting ratios to the *LR* of the data. We use the Benjamini-Hochberg procedure to correct for multiple comparisons, setting an FDR cutoff of 5%.

## Acknowledgments

We thank members of the Stuart lab for feedback on the methodology and Rocio Soto Astorga for graphics support. PC and JMS are supported by a grant from the California Institute of Regenerative Medicine working under the auspices of the Stem Cell Genome Center of Excellence. JMS was also supported by NIGMS grant 5R01GM109031.

## Method availability and supplementary material

SCIMITAR code and documentation are freely available at https://github.com/dimenwarper/scimitar. Supplementary methods and results can be found at https://github.com/dimenwarper/scimitar/wiki.

